# Multiplexed Molecularly-Specific SWCNT Cytokine Sensing is Enabled and Enhanced by Amine-Functionalized DNA Aqueous Two-Phase Extraction

**DOI:** 10.1101/2025.05.16.654499

**Authors:** Amelia Ryan, Sadiyah Parveen, Zachary Cohen, Atara Israel, Ryan Williams

**Affiliations:** The City College of New York, Department of Biomedical Engineering, New York, NY 10031

**Keywords:** Single-walled carbon nanotube, chirality, multiplex, sensors, inflammation

## Abstract

Single-walled carbon nanotubes (SWCNT) are versatile building blocks for optical sensors. Their carbon lattice structure, denoted by chiral (n,m) indices, determines the wavelength of their near infrared fluorescence emission. Commercially available SWCNT are polydisperse, containing many (n,m) species with overlapping fluorescence peaks, leading to spectral congestion. Aqueous two-phase extraction (ATPE) has emerged as a valuable tool to sort SWCNT according to their chiral structure and promises improved nanosensor performance as well as unique opportunities for multiplexing. Previous applications of ATPE have enabled ratiometric sensors, but simultaneous detection of multiple analytes using distinct SWCNT chiralities has yet to be realized. This hurdle is largely due to the contrasting surface chemistries needed for ATPE sorting and those needed for biosensing applications. In this work, we introduce a method for purifying multiple chiralities of SWCNT which can be functionalized with molecularly specific antibodies to create a multiplexed sensor platform. We utilize aminated DNA (DNA-NH_2_), which serves as both a conduit for ATPE sorting and an intermediate linker for later antibody conjugation, to realize this goal. We demonstrate that chemically modified DNA enables ATPE sorting, supporting the broad future utility of the technique. We further show that chirality-sorted SWCNT-DNA-NH_2_ serve as excellent base constructs for antibody conjugation, forming highly specific sensors with enhanced optical properties (compared to unsorted SWCNT sensors). Importantly, one batch of monochiral base constructs can be functionalized with different antibodies towards different biological applications. Finally, we show that we can sort and functionalize two separate SWCNT chiralities and combine them together to create a multiplexed sensor platform where each peak responds distinctly to its target. There are vast opportunities for expansion of this work through the use of alternative DNA sequences to enable sorting, various SWCNT chiralities, and virtually any commercially available antibody for molecular recognition.

## Introduction

Single-walled carbon nanotubes (SWCNT) are highly effective transducers for optical sensors. They emit near infrared (NIR) narrow-bandgap photoluminescence dependent on the chiral angle of the sp^2^ carbon lattice helix, a feature conducive to biomedical sensing applications.^1^ Each species is denoted by an (*n,m*) index defined by this chiral angle and tube diameter.

Though SWCNT possess such useful inherent properties, some amount of processing is necessary for these applications. As-produced SWCNT are hydrophobic and thus readily aggregate.^2^ Non-covalent functionalization with DNA is a widely used for solubilizing SWCNT as it disperses individual nanotubes in aqueous media due through π-π base stacking on the SWCNT surface and charge repulsion of the phosphate backbone.^3-5^ Such non-covalent solubilization allows for the bright, stable photoluminescence of SWCNT in solution, as well as imparting biocompatibility.^6, 7^ Long-term *in vivo* studies have shown that SWCNT wrapped with DNA show no signs of short-term or chronic toxicity.^6^

Synthesis of SWCNT-DNA constructs has allowed for further screening and development, with or without the addition of biomolecular recognition elements, as effective biosensors. Many studies have utilized SWCNT-DNA to detect a wide variety of analytes: small molecules such as H_2_O_2_, and doxorubicin, neurotransmitters including dopamine, norepinephrine, and serotonin, and protein biomarkers ranging from gynecologic cancer-associated proteins to inflammatory cytokines.^8-13^ Several studies have used functionalized DNA oligonucleotides to anchor molecularly specific probes to the SWCNT surface without compromising the optical properties of the SWCNT. Other studies have used DNA as a base to conjugate further sensor elements. In particular, primary amine-functionalized DNA (SWCNT-DNA-NH_2_) is necessary to attach an antibody to the nanotube construct.^14, 15^ Carbodiimide crosslinking has been used to form an amide bond between the DNA-NH_2_ and the antibody, creating a stable bond and molecular-specific nanobiosensor. This antibody-based approach has been used to target ovarian cancer biomarker HE4, prostate cancer biomarker uPA, the estrogen receptor, and inflammatory cytokine interleukin-6 (IL-6) with high specificity.^14-17^

Commercially available SWCNT are comprised of many unique (*n,m*) species, and thus have many, often-overlapping, optical transitions.^1, 18, 19^ Circumventing such spectral overlap in SWCNT transducer NIR photoluminescence is necessary to produce discrete, highly sensitive sensors, as well as to produce highly-multiplexed sensors.^20^ SWCNT chiral sorting and purification is an enabling step towards these goals.^21^ Several effective separation techniques developed over the past two decades include density gradient centrifugation, gel chromatography, ion exchange chromatography, and aqueous two-phase extraction (ATPE).^18, 22-24^

ATPE is widely-used for nanomaterials separation in general, and SWCNT separation specifically, as it is versatile, scalable, and relatively low-cost.^18, 25-34^ Some ATPE separations used bile salt surfactants to solubilize SWCNT prior to sorting and subsequent surfactant exchange with DNA.^26, 35-37^ Others used DNA to solubilize SWCNT prior to sorting that was subsequently used directly in sensing applications.^38-41^ This is analogous to prior ion-exchange chromatography separation methods, which relied on ssDNA recognition of individual (*n,m*) species.^24^ However, this sorting method has thus far limited chirally-pure SWCNT sensor development to analytes that can be detected with the same sequence used for chirality sorting. Further functionalization of SWCNT-DNA sorted through ATPE, such as the above-mentioned carbodiimide cross-linking with antibodies has yet to be realized.

In this work, we demonstrated ATPE sorting of SWCNT with primary amine-functionalized DNA, which enabled subsequent molecularly-specific multiplexed sensing for the first time. We evaluated separation patterns, side-by-side, of both non-functionalized and aminated versions of several known ATPE-active DNA sequences. We found that the ATPE method is indeed robust, with some pattern differentiation, to functional and non-functional DNA sequence incorporation. We then performed antibody conjugation directly to the purified SWCNT-DNA-NH_2_, producing the first multiplexed SWCNT sensors functionalized with commercial biomolecular recognition elements—in this case for the inflammatory cytokines IL-6 and IL-12. These targets were chosen due to their crucial roles in cell-mediated immunity. IL-6 and IL-12 regulate both acute and chronic inflammation and are valuable biomarkers of inflammatory diseases.^42, 43^ The framework presented here could be applied to a wide variety of targets as the vast library of commercially available antibodies only broadens the utility of this straightforward multiplexed sensor construction method.

## Materials and Methods

### Preparation of SWCNT-DNA Constructs

SWCNT-DNA was prepared as previously described.^12^ The (6,5) sorting sequence (ss65, TTA-TAT-TAT-ATT) and (7,6) sorting sequence (ss76, (ATTT)_4_) were identified as chirality-recognizing sequences in previous ATPE studies.^28,31^ Amine-functionalized versions of each sequence were also used. (TAT)_6_-NH_2_ was used as an intermediate linker for antibody conjugation on unsorted SWCNT as previously published^14-16^ (**Table 1**). 1 mg DNA (Integrated DNA Technologies; Coralville, IA) as added to 0.5 mg SWCNT powder (SG65i [Sigma-Aldrich; Burlington, MA], SG76 [Sigma-Aldrich; Burlington, MA], or HiPCO [Nanointegris; Boisbriand, Quebec]) in 0.5 mL deionized water with 0.1 M NaCl. Samples were sonicated on ice at 40% amplitude for 1 h by a VCX 750 ultrasonicator with a 2mm stepped microtip probe (Sonics & Materials, Inc.; Newtown, CT). Sonicated suspensions were ultracentrifuged at 58,000 Xg for 1 h using an Optima Max-XP Ultracentrifuge (Beckman Coulter; Brea, CA). The top 75% of the suspension was collected for further use. Samples were stored at 4°C for up to 14 days. Within 24 h of use, samples were filtered to remove free DNA. An aliquot of SWCNT-DNA was loaded into a 100 kDa MWCO centrifugal filter (Sigma-Aldrich; Burlington, MA) and centrifuged for 15 min at 14,000 Xg. Filtrate was discarded, and the contents of the filter were resuspended in deionized water, then centrifugally filtered again. The retained content was collected and resuspended in deionized water for further analysis.

**Table 1.** ssDNA Sequences Used for SWCNT Dispersion.

| Name | Sequence |
| --- | --- |
| ss65 : (6,5) sorting sequence<sup>31</sup> | TTA-TAT-TAT-ATT |
| ss65-NH<sub>2</sub>: aminated (6,5) sorting sequence | TTA-TAT-TAT-ATT-NH<sub>2</sub> |
| ss76 : (7,6) sorting sequence<sup>44</sup> | (ATTT)<sub>4</sub> |
| ss76-NH<sub>2</sub>: aminated (7,6) sorting sequence | (ATTT)<sub>4</sub>-NH<sub>2</sub> |
| (TAT)<sub>6</sub>-NH<sub>2</sub>: intermediate linker sequence<sup>14-17</sup> | (TAT)<sub>6</sub>-NH<sub>2</sub> |

### Synthesis of ATPE Systems and Mimics

ATPE systems were prepared as described by others.^31^ 400 μL ATPE systems were made with 57 μL of 60% (m/m) 1.5kDa polyethylene glycol (PEG) (Sigma-Aldrich; Burlington, MA), 202 μL of 20% (m/m) 250 kDa Dextran (DEX) (Alfa Aesar; Haverhill, MA), 21 μL of deionized water, and 120 μL of ∼ 500 mg/L SWCNT-DNA or SWCNT-DNA-NH_2_. As ATPE is a scalable technique, 3200 μL systems were prepared using the same ratio of reagents with volumes multiplied 8x.^18^

Similarly, the ATPE mimic used the same amount of PEG and DEX but instead replaced the SWCNT dispersion with the identical volume of water. For our experiments, we chose to scale the mimic 30x from the ATPE system in order to have sufficient polymer volume for our separations. The mimic was vortexed for 1 minute and centrifuged at room temperature for 5 mins at 3,200 Xg to allow for phase (PEG/DEX) separation.

### ATPE Separation of SWCNT

2 μL of 1% polyvinylpyrrolidone (PVP, 10 kDa, Sigma-Aldrich; Burlington, MA) was initially added to the 400 μL ATPE system. We vortexed the system for 1 minute and centrifuged at 3,200 Xg for 5 mins or until a clear phase separation was visible. After centrifugation, the top fraction of the ATPE system was extracted via micropipette and saved for optical characterization. The extracted top phase was replaced with an equal volume of the top phase of the mimic solution to keep total volume and polymer concentrations consistent throughout the chirality sorting process. 2 μL 1.25% PVP was added along with the mimic. The ATPE system was then vortexed and centrifuged as before. The top phase was again extracted and saved for optical characterization. This process was repeated, with the addition of 2 μL 1.25% PVP each iteration, until the purified target chirality was present, whether it be in the top or bottom phase. Only when sorting with ss76 or ss76-NH_2_ was 2 μL 2% PVP used starting with the 4^th^ extraction and continuing until the end.

### Optical Characterization of SWCNT

At each step of the ATPE process, SWCNT fractions were subjected to absorbance measurements (V-730 UV–vis Spectrophotometer, Jasco; Easton, MD) from 300–1100 nm using a with a 400 nm min^−1^ scan rate and 0.2 nm steps. The concentration of SWCNT was determined using the equation: C = A_630_/0.036 l mg^−1^ cm^−1^ * DF, where DF = dilution factor.^45^ We also performed NIR fluorescence spectroscopy on each SWCNT fraction using a NS MiniTracer (Applied NanoFluorescence; Houston, TX) using a 50 mW 638 nm laser in the range of 900–1600 nm. After optical characterization was complete, samples were kept in the 4°C fridge until use.

### Engineering Molecularly-Specific Sensors

We then conjugated cytokine-specific antibodies to the chirally-sorted SWCNT-DNA-NH_2_ samples via carbodiimide conjugation chemistry.^14-16^ Separately, as a non-separated comparison, antibody conjugation was also performed on HiPCO (bulk chirality) SWCNT wrapped with (TAT)_6_-NH_2_. IL-6 (RRID: AB_398568, BD Biosciences; Franklin Lakes, NJ) and IL-12 (RRID: AB_2929120, PeproTech; Cranbury, NJ) antibody conjugations were performed separately. The carboxylic acids of the antibody were first activated with 1-ethyl-3-(3-dimethylainopropyl) carbodiimide (Fisher Scientific; Hampton, NH) and N-hydroxysuccinimide (Fisher Scientific; Hampton, NH) in a 10X and 25X molar excess, respectively, for 15 min. This reaction was quenched with 1 μL of 2-mercaptoethanol (Fisher Scientific; Hampton, NH). The activated antibody was added in an equimolar ratio to the DNA, assuming a 1:1 ratio of ssDNA to SWCNT. Following 2 hours of incubation on ice, the conjugate was dialyzed against water with a 1,000 kDa MWCO SpectraPor Float-A-Lyzer (Repligen; Waltham, MA) at 4°C for 48 hours with two buffer changes to remove unconjugated antibody, reaction reagents, and leftover ATPE polymers from the solution. The resulting sensors – made from IL-6 antibody conjugated to (7,6)-enriched SWCNT (IL-6 Ab-(7,6)) and IL-12 antibody conjugated to (6,5)-enriched SWCNT (IL-12 Ab-(6,5)) – were collected from dialysis and stored at 4°C. After sample collection, we performed light scattering measurements to confirm successful antibody conjugation. Dynamic light scattering was used to measure the particle size before and after conjugation while electrophoretic light scattering was performed to compare the relative zeta potential of the particles before and after conjugation (Nano-ZS90, Malvern; Worcestershire, U.K.).

### Sensor Characterization

Prior to use, sensors were passivated by incubation with 50x mass ratio poly-L-lysine (PLK, Advanced BioMatrix; Carlsbad, CA) for 30 mins at 4°C, as previously-described to direct analyte binding to the antibody rather than the exposed SWCNT surface.^16^ We incubated 0.5 mg/L SWCNT sensors with 5 μg/mL of the target cytokine protein (Human IL-6 Recombinant Protein, Gibco; Waltham, MA **or** Recombinant Human IL-12 (linked heterodimer), R&D Systems; Minneapolis, MN) in 1X PBS to confirm functionality. NIR fluorescence spectra were obtained with ClaIR plate reader (Photon Etc; Montreal, Quebec) in 15-minute intervals for 3 hours following protein addition.

To evaluate multiplexing ability, an equal mass of IL-6 Ab-(7,6) and IL-12 Ab-(6,5) sensors were combined into one sample. The combined sensors were tested against IL-12 individually, IL-6 individually, as well as both cytokines combined. The specificity of the sensors was further challenged by testing them against non-target cytokines IL-1β and TNF-α. Sensor dynamic range was investigated using a broad range of analyte concentrations (from 15 pg/mL to 10 μg/mL). All sensor samples were prepared in triplicate unless otherwise specified.

### Data Analysis

NIR fluorescence corresponding to individual nanotube chirality emission peaks was fit with a custom MATLAB code to a Voigt model to determine their center wavelength and maximum intensity values (MATLAB code is available upon request). Changes in the SWCNT center wavelength and maximum intensity values were reported relative to their baseline emission prior to antigen addition. Means for triplicate samples plus the standard deviation were obtained. Statistical significance was determined with a two-sample *t* test.

## Results and Discussion

### ATPE Sorting @of SWCNT-DNA-NH_2_

The (6,5) Super Sequence, or ss65, is a known recognition sequence for (6,5) SWCNT.^31^ As expected, it enabled purification of the (6,5) chirality from SG65i-prepared SWCNT stock (**Figure 1A, 1B, S1A)**, wherein the T2 Fraction demonstrates strong (6,5) purification. However, separation of SWCNT with NH_2_-functionalized ss65 exhibited markedly different behavior while still being able to achieve separation. (6,5) nanotubes encapsulated with ss65-NH_2_ appear more resistant to partitioning to the top phase in that they require more PVP to move to the top phase in comparison to their non-aminated counterparts (**Figure 1C, S1B**). Although it required a higher number of extractions (and more PVP) to achieve the same purity, ss65-NH_2_ also enabled (6,5) purification in the B4 and T5 fractions.

**Figure 1.**
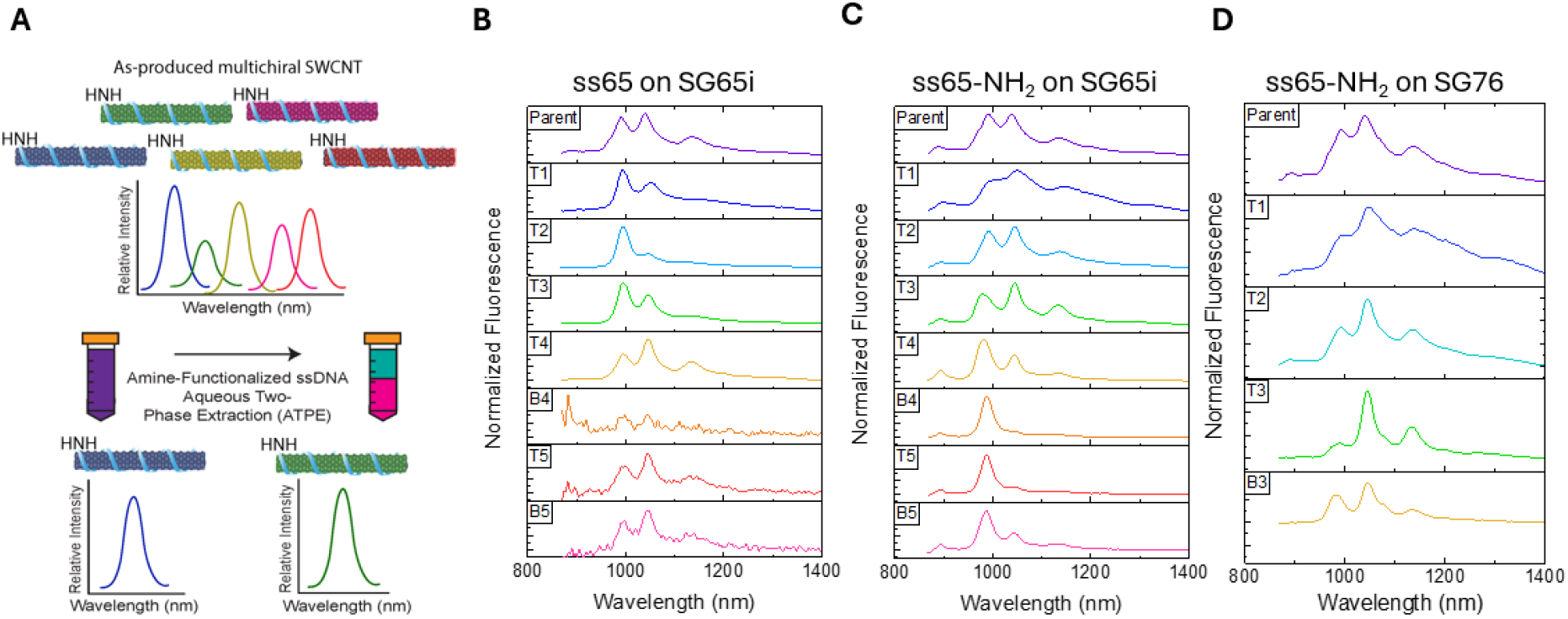
ATPE sorting with ss65-NH_2_. **A)** Diagram of workflow. **B)** Fluorescence spectra acquired with 638 nm laser from each phase of ATPE sorting SG65i SWCNT using ss65. **C)** Fluorescence spectra from each phase of sorting SG65i SWCNT using ss65-NH_2_. **D)** Fluorescence spectra from each phase of sorting SG76 SWCNT using ss65-NH_2_.

One potential explanation for this observation is differences in hydrophilicity of the SWCNT-ss65 and the SWCNT-ss65-NH_2_. The amine group contains a nitrogen atom with a lone pair of electrons that can form hydrogen bonds with water molecules – increasing the hydrophilic character of the DNA strand.^46^ We hypothesize that this may increase the overall hydrophilicity of the resulting SWCNT constructs, in turn making each (*n,m*) species more likely to partition to the hydrophilic bottom phase of the ATPE system.^31^ Therefore, more PVP partitioning modulator is needed to extract the SWCNT-DNA-NH_2_ from the bottom phase compared to the non-aminated SWCNT-DNA.^18, 28, 31^ This difference in hydrophilicity may enable modified and potentially even enhanced ATPE sorting resolution. It also suggests that functional groups on DNA that affect the SWCNT construct’s solvation energy, such as a primary amine group, could be used to fine-tune ATPE resolution. This is the first instance of chemically-modified DNA enabling ATPE sorting of SWCNT. It demonstrates the robustness of the ATPE method, but also the possibility for further customization of chirality-sorted SWCNT through the use of DNA with other functional groups.

We further assessed the ability of amine-functionalized DNA to enable SWCNT chirality sorting using SG76-prepared SWCNT, also using the ss65-NH_2_ sequence. In this study, iterative additions of PVP to the ATPE system enabled a (7,5)-enriched fraction to emerge in T3 (**Figure 1D, S1C**). These results suggest that the robustness of the ATPE method is preserved when using amine-functionalized DNA to sort multiple chiralities. Although these experiments yielded chirality-enriched fractions of SWCNT-DNA-NH_2_, the concentration of SWCNT in the final products was relatively low. Thus, we sought to assess DNA-NH_2_-enabled sorting scale-up as previously demonstrated with ATPE.^31, 34, 47^

In order to achieve sufficient quantities of purified SWCNT-DNA-NH_2_, we scaled up the separation by a factor of eight. We observed similar purities of (6,5) achieved in the scaled sorting system (**Figure S2A, S3A**). We also investigated the use of ss76 as it has been reported as a (7,5) recognition sequence in ion exchange chromatography.^28^ However, when used in the ATPE sorting system, ss76-NH_2_ enabled gradual extraction of almost all chiralities except (7,6) from SG76 nanotubes, leaving purified (7,6) in the bottom phase (**Figure S2B, S3B**). While we found minor variations between the results of the 400 μL and 3200 μL systems, the sorting behavior of the scaled-up system was largely predictable. For our applications, we prioritized high-yield samples, using (6,5)-enriched fractions that contain no (7,6) and (7,6)-enriched fractions that contain no (6,5), rather than completely monochiral samples with lower yield. The scaled-up system yielded sufficient amounts of purified SWCNT for our antibody conjugation process, though we anticipate that larger scales could be used for future applications.

### Antibody Conjugation

To demonstrate the potential of multiplexed SWCNT-based sensors, we incorporated a standard antibody conjugation methodology with the chirality-enriched SWCNT. We conjugated an anti-IL-12 antibody to (6,5)-enriched SWCNT to produce IL-12 Ab-(6,5) and anti-IL-6 to the (7,6)-enriched SWCNT to produce IL-6 Ab-(7,6). Optical characterization revealed that the SWCNT constructs retained their fluorescence signal after antibody conjugation (**Figure 2A**). When the sensors were combined, both the (6,5) and (7,6) chiralities show pronounced peaks, demonstrating that these SWCNT can be used for spectral multiplexing without overlap.

**Figure 2.**
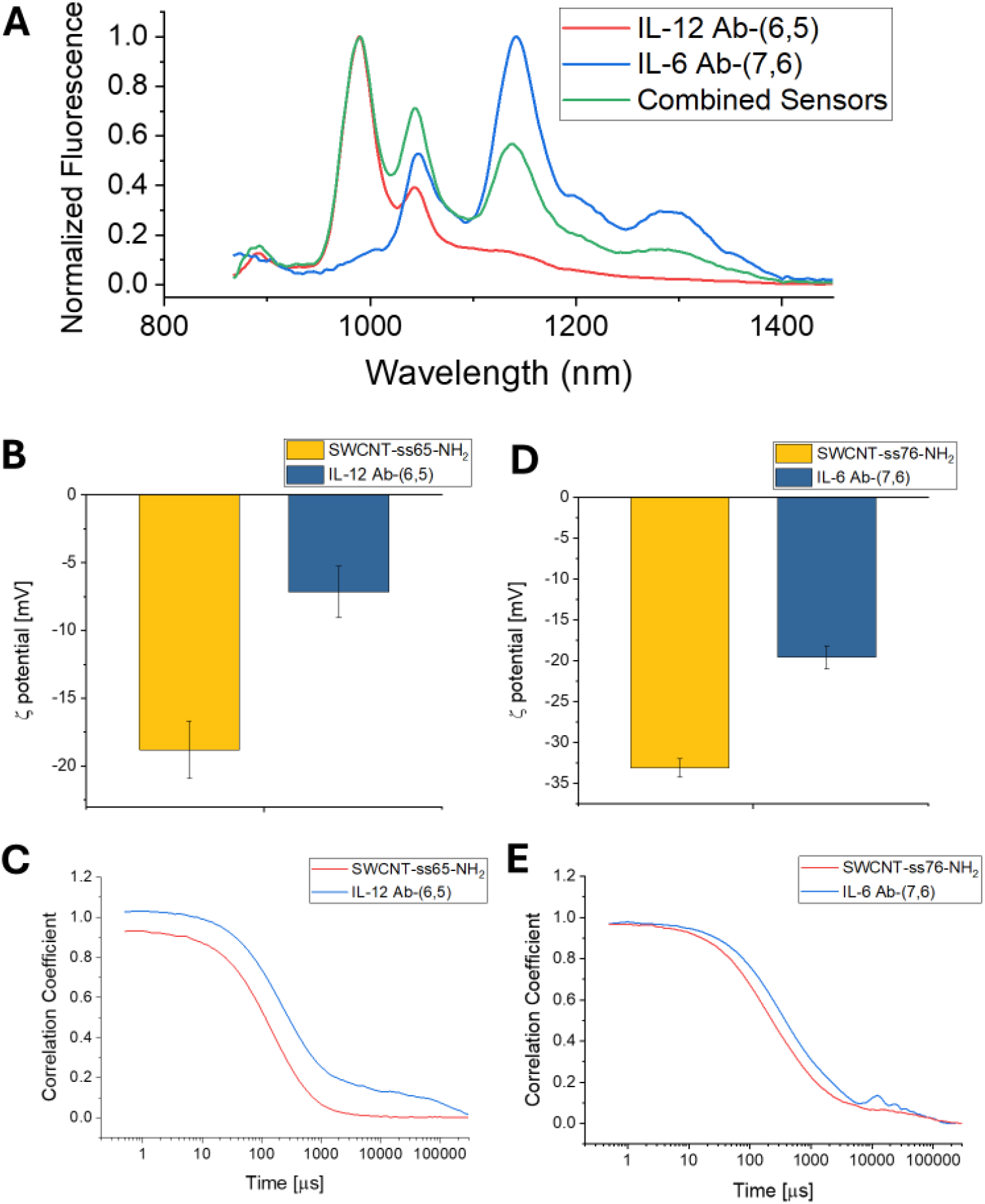
Optical characterization of antibody-conjugated nanosensors. **A)** Representative fluorescence spectra acquired with 638 nm laser of antibody-conjugated nanosensors before deployment. **B)** Electrophoretic light scattering indicates (6,5) SWCNT construct increased in charge after IL-12 antibody conjugation. **C)** Electrophoretic light scattering indicates (7,6) SWCNT construct increased in charge after IL-6 antibody conjugation. **D)** Dynamic light scattering indicates (6,5) SWCNT construct increased in size after IL-12 antibody conjugation. **E)** Dynamic light scattering indicates (7,6) SWCNT construct increased in size after IL-6 antibody conjugation.

Although there is a considerable (7,5) peak in the fluorescence measurement of our combined sensors, we found that this did not inhibit the function of our multiplexed sensor system. Further sensor characterization confirmed successful antibody conjugation as the relative size and ζ-potential of the SWCNT constructs increased after conjugation (**Figure 2B-2E)**.

There have been several examples of carbodiimide crosslinking performed to attach antibodies to aminated DNA on SWCNT for optical sensing, all of which have used unsorted SWCNT and have exclusively used (TAT)_6_-NH_2_ as the chosen DNA sequence.^14-17^ Our successful conjugation of antibodies to ATPE-sorted fractions in this work establishes that chirality-sorted SWCNT can also serve as base constructs for antibody-conjugated sensors. Furthermore, we have shown that the antibody conjugation reaction can be performed in environments containing PEG and DEX solutions. This work also demonstrates that aminated DNA sequences other than (TAT)_6_-NH_2_ can be used for this application.

Other works have utilized SWCNT-DNA for ATPE sorting and subsequent sensing with no further biomolecular recognition element functionalization.^38-41^ However, these sensing applications have been limited to analytes that can be detected using non-selective DNA sequences used for ATPE sorting. Other studies have circumvented this limitation by sorting surfactant-dispersed SWCNT, then switching out the surfactant for the DNA sequence of choice after ATPE is completed.^26, 35-37^ Our work here provides a more direct and versatile alternative: sorted SWCNT fractions from ATPE can be immediately functionalized with any antibody of choice. The vast library of antibodies allows for virtually any biomarker to be targeted. The direct route to functionalization improves efficiency and prevents loss of carbon material. These new developments open a myriad of possibilities in the world of SWCNT research.

### Multiplexed Sensing with Chirality-Enriched SWCNT

After chirality purification and antibody conjugation, we anticipated that IL-12 Ab-(6,5) would enable a shift in the (6,5) peak in response to IL-12 protein (**Figure 3A**). When tested individually, IL-12 Ab-(6,5) exhibited an average red shift of 1.76 nm +/-0.16 nm after 3 hours of exposure to 5 μg/mL IL-12 protein (**Figure 3B**). After confirming the basic functionality of IL-12 Ab-(6,5), we combined it with IL-6 Ab-(7,6) to assess simultaneous function. IL-12 Ab-(6,5) exhibited a red shift in response to samples containing IL-12 only and IL-12 in addition to IL-6 (**Figure 3C, S4, S5**). Importantly, the (6,5) peak showed a significant red shift in response to IL-12 protein, while the (7,6) peak did not (**Figure 3D, 3E**). These results demonstrate the ability of the monochiral analyte-specific sensors to function independently of each other.

**Figure 3.**
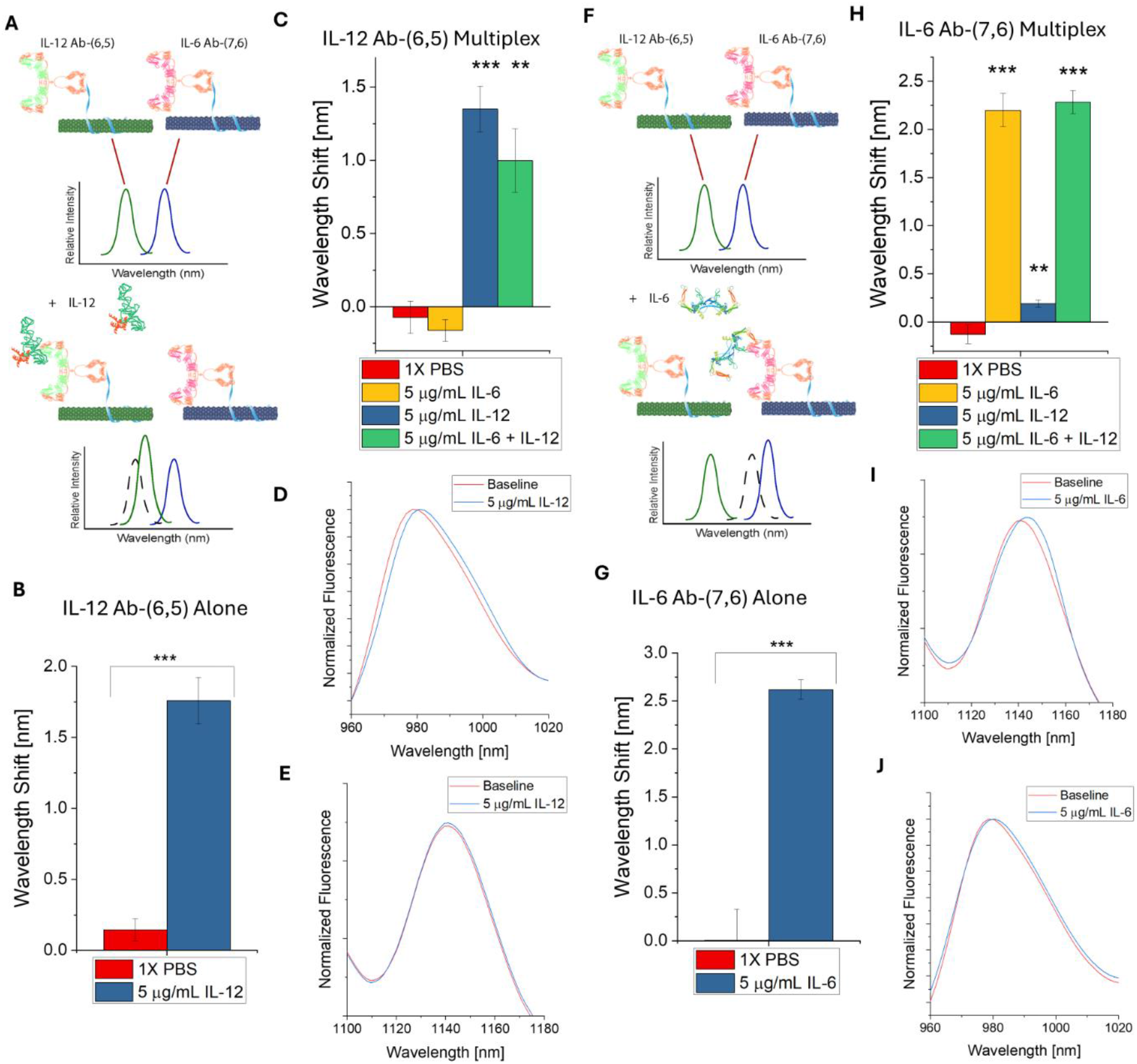
Multiplexed cytokine detection using chirality-sorted SWCNT-antibody sensors. **A)** Schematic of anticipated (6,5) peak response to IL-12. **B)** Wavelength shift of IL-12 Ab-(6,5) alone in response to 5 μg/mL IL-12 protein. **C)** Wavelength shift of (6,5) peak in combined sensor samples response to cytokine addition (acquired with 655 nm laser). **D)** Representative (6,5) spectral shift from combined sensors in response to IL-12 only. **E)** Representative (7,6) spectral shift from combined sensors in response to IL-12 only. **F)** Schematic of anticipated (7,6) peak response to IL-6. **G)** Wavelength shift of IL-6 Ab-(7,6) alone in response to 5 μg/mL IL-6 protein. **H)** Wavelength shift of (7,6) peak in combined sensor samples response to cytokine addition. **I)** Representative (7,6) spectral shift from combined sensors in response to IL-6 only. **J)** Representative (7,6) spectral shift from combined sensors in response to IL-12 only.

Similarly, we anticipated that IL-6 Ab-(7,6) would enable a shift in the (7,6) peak in response to IL-6 protein (**Figure 3F**). When tested individually, IL-6 Ab-(7,6) exhibited an average red shift of 2.62 nm +/-0.10 nm in response to 5 μg/mL IL-6 protein (**Figure 3G**). When both sensors were combined, IL-6 Ab-(7,6) showed a robust red shift in response to samples containing IL-6 alone as well as IL-6 and IL-12 together (**Figure 3H, S4, S6**). As expected, the (7,6) peak shifts in response to IL-6 protein alone while the (6,5) peak does not (**Figure 3I, 3J**). In samples where both cytokines are present, both (6,5) and (7,6) peaks show a red shift (**Figure S4, S7**).

Here, we demonstrate spectral multiplexing using two different SWCNT chiralities in the same solution for the first time. The use of ATPE-purified SWCNT allows us to clearly interpret the optical signal of each chirality with minimal interference from other chiralities, metallic SWCNT, defective SWCNT, and other impurities. We have shown here that SWCNT of different chiralities can be sorted, functionalized separately against unique targets, and then combined into one sample to create a sensor array in which each chirality represents sensors against a different target.

We performed an additional challenge to the specificity of the sensors by testing them against inflammatory cytokines interleukin-1β (IL-1β) and tumor necrosis factor-α (TNF-α), two immunologically similar signaling molecules. Neither the (6,5) nor the (7,6) peak showed any significant shifts in response to IL-1β or TNF-α, further supporting the high molecular specificity of these sensor constructs (**Figure 4A, 4B**).

**Figure 4.**
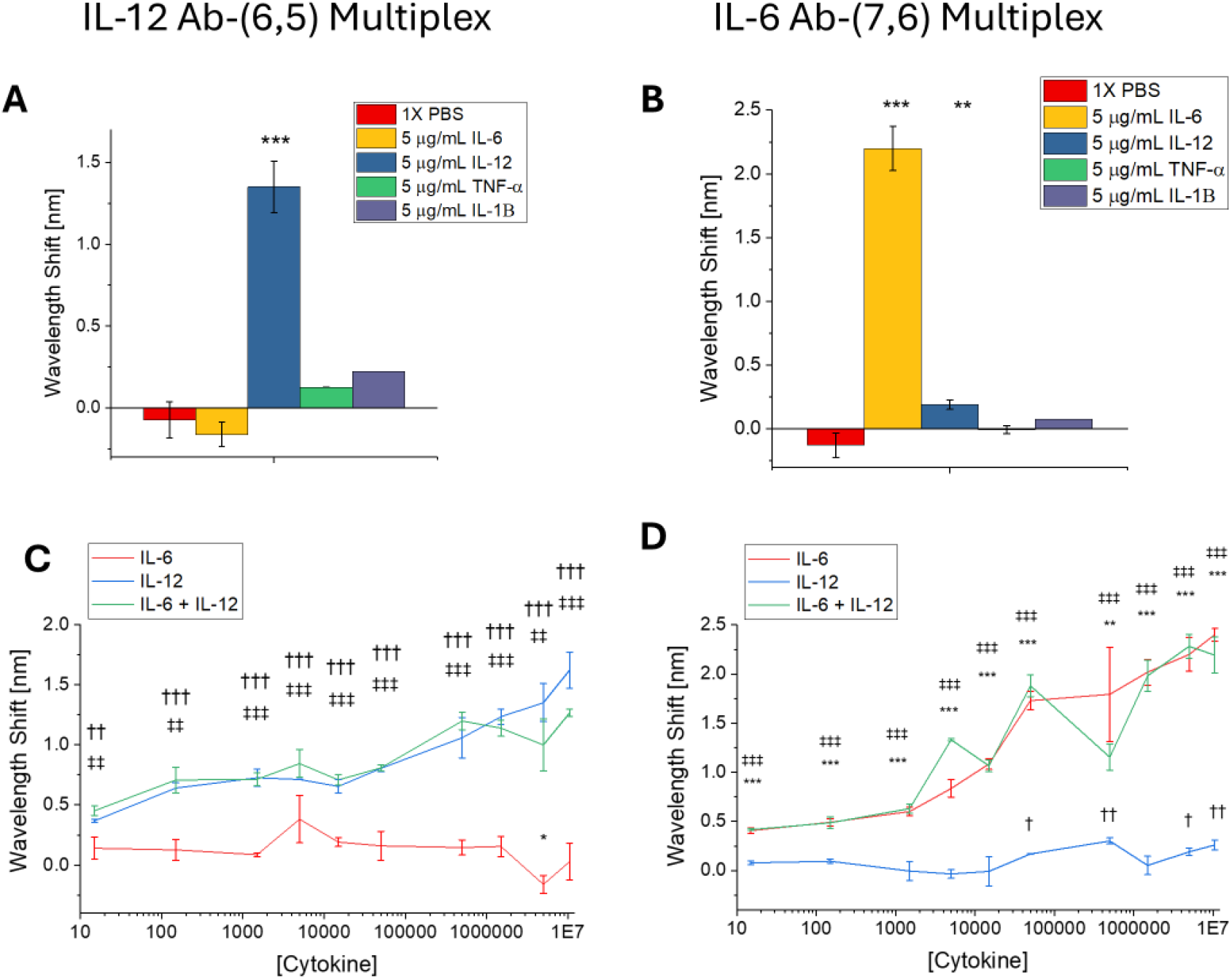
Selectivity and sensitivity of the multiplexed nanosensor. **A)** Wavelength shift of (6,5) peak in response to IL-6, IL-12, TNF-α (n = 2), and IL-1β (n = 1). **B)** Wavelength shift of (7,6) peak in response to IL-6, IL-12, TNF-α (n = 2), and IL-1β (n = 1). **C)** Concentration-response curve of the (6,5) peak (acquired with 655 nm laser). **D)** Concentration-response curve of the (7,6) peak.

We sought to further understand the functionality of the sensor array by testing its dynamic range and limit of detection (LOD). The sensors were again deployed against IL-12, IL-6, and both cytokines combined at concentrations from 10 μg/mL to 15 pg/mL, a wide range of physiologically relevant cytokine levels. Even when titrated to as low as 15 pg/mL, both (6,5) and (7,6) peaks showed significant wavelength shifts in response to their respective cytokine. The (6,5) peak exhibited 0.368 nm red shift (+/-0.013 nm) in response to 15 pg/mL IL-12. The (7,6) peak exhibited 0.407 nm red shift (+/-0.029 nm) in response to 15 pg/mL IL-6. Both peaks exhibited similar patterns of response to their target cytokine and the combined cytokine samples. 15 pg/mL IL-12 + IL-6 yielded 0.452 nm (+/-0.039 nm) from the (6,5) peak and 0.416 nm (+/ 0.0095 nm) from the (7,6) peak (**Table S1, S2**). Neither peak showed considerable shifts in response to their non-target cytokine alone (**Figure 4C, 4D)**. These results indicate that the response of the (6,5) peak is caused by the IL-12 antibody binding event, the response of the (7,6) peak is caused by the IL-6 antibody binding event, and that the sensor response is not additive.

The concentrations at which we demonstrated detection are well within the physiologically relevant ranges for both cytokines – healthy levels of IL-6 in human serum are reported from 0 to 181 pg/mL, while patients with inflammatory disease have elevated IL-6 serum levels. 567 pg/mL has been observed in Alzheimer’s disease patients, 692 pg/mL in rheumatoid arthritis patients, while extremely high levels – up to 500,000 pg/mL – are reported in cases of severe sepsis.^48-51^ Fewer studies have been published on clinically relevant levels of IL-12. Some report healthy patients do not show serum IL-12 levels higher than 5 pg/mL (the typical sensitivity limit of ELISA) while others cite healthy serum levels at 82 pg/mL. However, upper limits of IL-12 levels in serum have been reported as 164.3 pg/mL in major depressive disorder patients, 442.7 pg/mL in chronic hepatitis B patients, and 371.2 pg/mL in malaria patients.^49, 52-55^ We note that, in our study, detection as such small concentrations is enabled by the small variation in sensor center wavelength shifts, likely afforded by the uniformity of the separated SWCNT transducers used.

In our literature search, we found no previously published optical sensors and only two examples of electrochemical sensors for IL-12 with LODs of 500 pg/mL and 3.5 pg/mL.^56, 57^ Literature on IL-6 sensors is much more extensive. Previously published sensors for IL-6 show a wide range of detection limits depending on their modality. Electrochemical sensors can detect IL-6 in the low pg/mL range, with one electrochemical impedance sensor detecting IL-6 in serum at a low limit of 0.01 fg/mL.^50, 58^ Optical sensors use surface plasmon resonance, fluorescence, and surface-enhanced Raman scattering to achieve LODs as low as 1.1 pg/mL, 0.02 pg/mL, and 0.028 pg/mL, respectively.^59-62^ Previously published SWCNT optical sensors for IL-6 have reported LODs of 3.91 μg/mL and 105 ng/mL using corona phase molecular recognition and aptamer-based approaches, respectively.^12, 63^ Our group has reported detection as low as 25 pg/mL using the same antibody-based design as the present study, but with polydisperse SWCNT rather than chirality-enriched.^16^

We then sought to assess the contributions of chirality-purified SWCNT to sensor robustness and repeatability. We compared chirally-pure SWCNT sensors to bulk chirality SWCNT sensors using the same antibody recognition element. Multi-chiral HiPCO SWCNT functionalized with the anti-IL-12 exhibited a significant red shift in response to recombinant IL-12 protein for all chiralities analyzed, though the degree of shifting varies among chiralities (**Figure 5A**). The (6,5) peak is not visible in the HiPCO mixture at the laser excitation wavelength used, so this peak cannot be directly compared to IL-12 Ab-(6,5). The mean wavelength shift exhibited by the (9,5) chirality in the HiPCO sensors (1.79 nm) is similar to the mean shift exhibited by IL-12 Ab-(6,5) (1.76 nm) in the same testing conditions. However, the standard deviation of the chirality-sorted sensors is lower (0.244 nm in HiPCO sensors, 0.164 nm in (6,5) sorted sensors). There is also less shifting and smaller variation in the control group of IL-12 Ab-(6,5) compared to the unsorted HiPCO sensors, indicating less signal interference in the chirality-sorted SWCNT. Multi-chiral HiPCO SWNCT functionalized with anti-IL-6 were also tested against 5 μg/mL IL-6 protein in 1X PBS. The (7,5) and (7,6) chiralities did not exhibit significant wavelength shifts, while the (9,5) chirality showed a mean shift of 1.55 nm with a standard deviation of 0.73 nm (**Figure 5B**). The shift observed in (9,5) is both lower in magnitude and less statistically significant than the shift observed in IL-6 Ab-(7,6).

**Figure 5.**
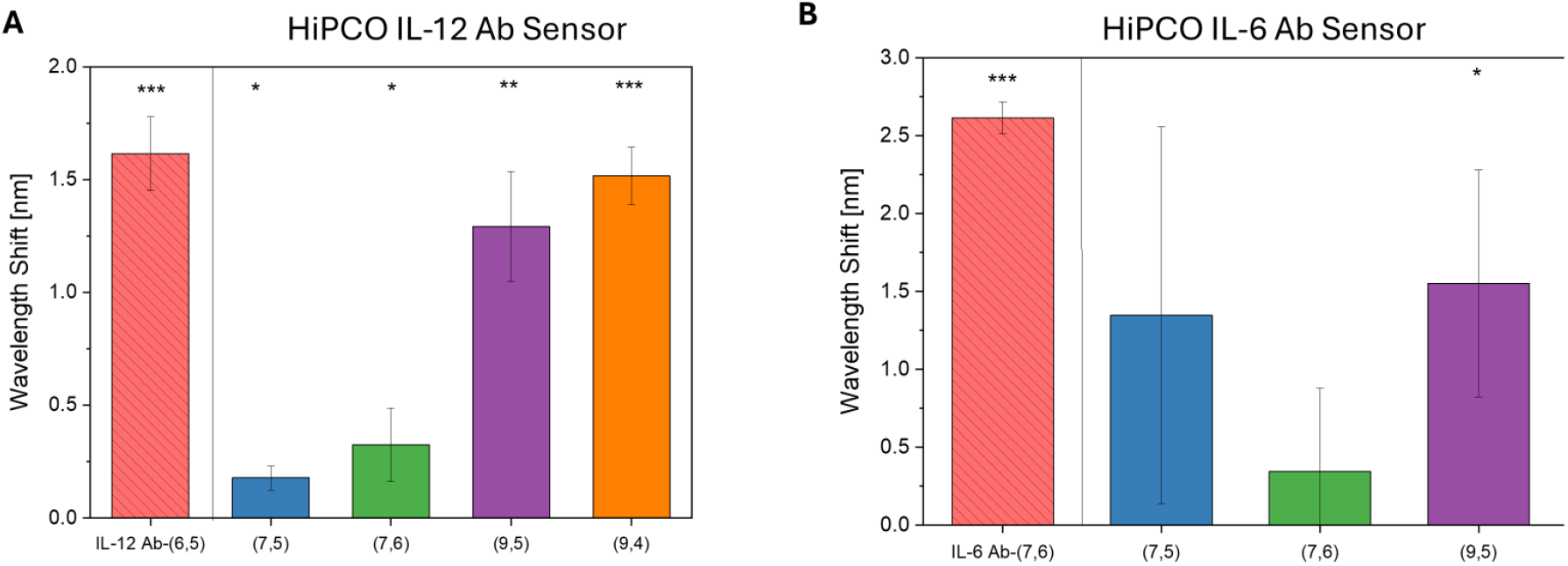
Chirally-pure sensor comparison to bulk chirality sensors. **A)** Wavelength shifts of IL-12 Ab-(6,5) and anti-IL-12-conjugated HiPCO SWCNT in response to 5 μg/mL IL-12 protein. (acquired with 638 nm laser) **B)** Wavelength shifts of IL-6 Ab-(7,6) and anti-IL-6-conjugated HiPCO SWCNT in response to 5 μg/mL IL-6 protein.

These results support the long-held hypothesis that chirality sorting of SWCNT improves optical sensor performance. The chirality-sorted sensors demonstrated less fluctuation in the center wavelength of the control groups, less variance among triplicates, and more robust statistical significance compared to polydisperse sensors made with the same antibodies. Thus, chiral purification will likely enhance the function of other SWCNT sensor designs as well and therefore holds great promise for the field of optical nanosensors.

### Conclusion and Future Work

In this work, we demonstrated ATPE sorting of (6,5) and (7,6) chiralities using amine-functionalized ssDNA, which is the first instance of chemically modified DNA enabling chirality purification. While the addition of the amine group altered the sorting behavior of the SWCNT within the ATPE system, our results suggest that aminated or otherwise functionalized DNA could be used to further customize sorting systems and subsequent applications of SWCNT obtained. We then performed antibody conjugation to two distinct SWCNT-DNA-NH_2_ constructs obtained from ATPE sorting, creating molecularly specific chirality-purified nanosensors. This unique sensor design combines the enhanced optical properties achieved by ATPE sorting, the high specificity enabled by antibodies, and, importantly, the opportunity for spectral multiplexing by functionalizing different (*n,m*) species with different analyte-specific recognition elements.

We are confident that the plug-and-play nature of the chirality-purified SWCNT-DNA-NH_2_ can serve as base constructs for virtually any antibody-based sensor. Future work will likely see this versatile approach utilized for other disease biomarkers, provided there is a suitable commercially available antibody to use for conjugation. The virtually unlimited library of DNA sequences and multitude of chemical modifications for oligonucleotides suggest the possibility for expansion of these methods. In addition, the application of machine learning to identify DNA sequences for chirality sorting may substantially improve the efficiency of the ATPE process.^33, 47^ Future studies may see the addition of aminated DNA sequences to machine learning training sets to identify more efficient sequences for sorting of SWCNT-DNA-NH_2,_ potentially coupled with molecular dynamics simulations to understand the functional group’s impact on SWCNT partitioning. Although (6,5) and (7,6) SWCNT were used in this work, this methodology could easily be expanded to additional chiralities, provided the appropriate DNA sequences are used to target different (*n,m*) species. Ultimately, we anticipate that this work will serve as a potential launchpad for translation of implantable or bedside multiplexed diagnostic tools.

## Acknowledgements

The authors greatly appreciate the insight and ATPE training of Ming Zheng of the National Institute of Standards and Technology. We also wish to acknowledge all members of the Williams Lab for discussion and feedback. This work was supported by NIH R35GM142833 (R. Williams) and The City College of New York Grove School of Engineering. A. Ryan and A. Israel were supported by a G-RISE Ph.D. traineeship from the National Institutes of Health (T32GM136499).

## References

(1) Bachilo, S. M.; Strano, M. S.; Kittrell, C.; Hauge, R. H.; Smalley, R. E.; Weisman, R. B. Structure-Assigned Optical Spectra of Single-Walled Carbon Nanotubes. Science 2002, 298, 2361–2366.

(2) O’Connell, M. J.; Bachilo, S. M.; Huffman, C. B.; Moore, V. C.; Strano, M. S.; Haroz, E. H.; Rialon, K. L.; Boul, P. J.; Noon, W. H.; Kittrell, C.; Ma, J.; Hauge, R. H.; Weisman, R. B.; Smalley, R. E. Band gap fluorescence from individual single-walled carbon nanotubes. Science 2002, 297 (5581), 593–596.

(3) Zheng, M.; Jagota, A.; Semke, E. D.; Diner, B. A.; McLean, R. S.; Lustig, S. R.; Richardson, R. E.; Tassi, N. G. DNA-assisted dispersion and separation of carbon nanotubes. Nat Mater 2003, 2 (5), 338–342.

(4) Jeng, E. S.; Moll, A. E.; Roy, A. C.; Gastala, J. B.; Strano, M. S. Detection of DNA Hybridization Using the Near-Infrared Band-Gap Fluorescence of Single-Walled Carbon Nanotubes. Nano Lett 2006, 6 (3), 371–375.

(5) Lee, A. J.; Wang, X.; Carlson, L. J.; Smyder, J. A.; Loesch, B.; Tu, X.; Zheng, M.; Krauss, T. D. Bright Fluorescence from Individual Single-Walled Carbon Nanotubes. Nano Lett 2011, 11 (4), 1636–1640.

(6) Galassi, T. V.; Antman-Passig, M.; Yaari, Z.; Jessurun, J.; Schwartz, R. E.; Heller, D. A. Long-term in vivo biocompatibility of single-walled carbon nanotubes. PLOS ONE 2020, 15 (5), e0226791.

(7) Kim, M.; Goerzen, D.; Jena, P. V.; Zeng, E.; Pasquali, M.; Meidl, R. A.; Heller, D. A. Human and environmental safety of carbon nanotubes across their life cycle. Nature Reviews Materials 2024, 9 (1), 63–81.

(8) Giraldo, J. P.; Landry, M. P.; Kwak, S.-Y.; Jain, R. M.; Wong, M. H.; Iverson, N. M.; Ben-Naim, M.; Strano, M. S. A Ratiometric Sensor Using Single Chirality Near-Infrared Fluorescent Carbon Nanotubes: Application to In Vivo Monitoring. Small 2015, 11 (32), 3973–3984.

(9) Cohen, Z.; Alpert, D. J.; Weisel, A. C.; Ryan, A.; Roach, A.; Rahman, S.; Gaikwad, P. V.; Nicoll, S. B.; Williams, R. M. Noninvasive Injectable Optical Nanosensor-Hydrogel Hybrids Detect Doxorubicin in Living Mice. Advanced Optical Materials 2024, 12 (17), 2303324.

(10) Kruss, S.; Landry, M. P.; Vander Ende, E.; Lima, B. M. A.; Reuel, N. F.; Zhang, J.; Nelson, J.; Mu, B.; Hilmer, A.; Strano, M. Neurotransmitter Detection Using Corona Phase Molecular Recognition on Fluorescent Single-Walled Carbon Nanotube Sensors. Journal of the American Chemical Society 2014, 136 (2), 713–724.

(11) Salem, D. P.; Landry, M. P.; Bisker, G.; Ahn, J.; Kruss, S.; Strano, M. S. Chirality dependent corona phase molecular recognition of DNA-wrapped carbon nanotubes. Carbon 2016, 97, 147–153.

(12) Ryan, A. K.; Rahman, S.; Williams, R. M. Optical Aptamer-Based Cytokine Nanosensor Detects Macrophage Activation by Bacterial Toxins. ACS Sens. 2024, 9 (7), 3697–3706.

(13) Yaari, Z.; Yang, Y.; Apfelbaum, E.; Cupo, C.; Settle, A. H.; Cullen, Q.; Cai, W.; Roche, K. L.; Levine, D. A.; Fleisher, M.; Ramanathan, L.; Zheng, M.; Jagota, A.; Heller, D. A. A perception-based nanosensor platform to detect cancer biomarkers. Science Advances 2021, 7 (47), eabj0852.

(14) Williams, R. M.; Lee, C.; Heller, D. A. A Fluorescent Carbon Nanotube Sensor Detects the Metastatic Prostate Cancer Biomarker uPA. ACS Sens. 2018, 3 (9), 1838–1845.

(15) Williams, R. M.; Lee, C.; Galassi, T. V.; Harvey, J. D.; Leicher, R.; Sirenko, M.; Dorso, M. A.; Shah, J.; Olvera, N.; Dao, F.; Levine, D. A.; Heller, D. A. Noninvasive ovarian cancer biomarker detection via an optical nanosensor implant. Science Advances 2018, 4 (4), eaaq1090.

(16) Gaikwad, P.; Rahman, N.; Parikh, R.; Crespo, J.; Cohen, Z.; Williams, R. M. Optical Nanosensor Passivation Enables Highly Sensitive Detection of the Inflammatory Cytokine Interleukin-6. ACS Appl Mater Interfaces 2024, 16 (21), 27102–27113.

(17) Gaikwad, P. V.; Rahman, N.; Ghosh, P.; Ng, D. L.; Williams, R. M. Detection of estrogen receptor status in breast cancer cytology samples by an optical nanosensor. Advanced NanoBiomed Research 2025, 5 (1), 2400099.

(18) Khripin, C. Y.; Fagan, J. A.; Zheng, M. Spontaneous Partition of Carbon Nanotubes in Polymer-Modified Aqueous Phases. Journal of the American Chemical Society 2013, 135 (18), 6822–6825.

(19) Kruss, S.; Hilmer, A. J.; Zhang, J.; Reuel, N. F.; Mu, B.; Strano, M. S. Carbon nanotubes as optical biomedical sensors. Advanced Drug Delivery Reviews 2013, 65 (15), 1933–1950.

(20) Zheng, M.; Jagota, A.; Strano, M. S.; Santos, A. P.; Barone, P.; Chou, S. G.; Diner, B. A.; Dresselhaus, M. S.; Mclean, R. S.; Onoa, G. B.; Samsonidze, G. G.; Semke, E. D.; Usrey, M.; Walls, D. J. Structure-Based Carbon Nanotube Sorting by Sequence-Dependent DNA Assembly. Science 2003, 302 (5650), 1545–1548.

(21) Nißler, R.; Ackermann, J.; Ma, C.; Kruss, S. Prospects of fluorescent single-chirality carbon nanotube-based biosensors. Analytical Chemistry 2022, 94 (28), 9941–9951.

(22) Arnold, M. S.; Green, A. A.; Hulvat, J. F.; Stupp, S. I.; Hersam, M. C. Sorting carbon nanotubes by electronic structure using density differentiation. Nature Nanotechnology 2006, 1 (1), 60–65.

(23) Liu, H.; Tanaka, T.; Kataura, H. Optical Isomer Separation of Single-Chirality Carbon Nanotubes Using Gel Column Chromatography. Nano Lett 2014, 14 (11), 6237–6243.

(24) Lustig, S. R.; Jagota, A.; Khripin, C.; Zheng, M. Theory of Structure-Based Carbon Nanotube Separations by Ion-Exchange Chromatography of DNA/CNT Hybrids. The Journal of Physical Chemistry B 2005, 109 (7), 2559–2566.

(25) Nadeem, A.; Kindopp, A.; Wyllie, I.; Hubert, L.; Joubert, J.; Lucente, S.; Randall, E.; Jena, P. V.; Roxbury, D. Enhancing Intracellular Optical Performance and Stability of Engineered Nanomaterials via Aqueous Two-Phase Purification. Nano Lett 2023.

(26) Nißler, R.; Kurth, L.; Li, H.; Spreinat, A.; Kuhlemann, I.; Flavel, B. S.; Kruss, S. Sensing with Chirality-Pure Near-Infrared Fluorescent Carbon Nanotubes. Analytical Chemistry 2021, 93 (16), 6446–6455.

(27) Fagan, J. A.; Khripin, C. Y.; Silvera Batista, C. A.; Simpson, J. R.; Hároz, E. H.; Hight Walker, A. R.; Zheng, M. Isolation of Specific Small-Diameter Single-Wall Carbon Nanotube Species via Aqueous Two-Phase Extraction. Adv Mater 2014, 26 (18), 2800–2804.

(28) Ao, G.; Khripin, C. Y.; Zheng, M. DNA-Controlled Partition of Carbon Nanotubes in Polymer Aqueous Two-Phase Systems. Journal of the American Chemical Society 2014, 136 (29), 10383–10392.

(29) Fagan, J. A.; Hároz, E. H.; Ihly, R.; Gui, H.; Blackburn, J. L.; Simpson, J. R.; Lam, S.; Hight Walker, A. R.; Doorn, S. K.; Zheng, M. Isolation of >1 nm Diameter Single-Wall Carbon Nanotube Species Using Aqueous Two-Phase Extraction. ACS Nano 2015, 9 (5), 5377–5390.

(30) Ao, G.; Streit, J. K.; Fagan, J. A.; Zheng, M. Differentiating Left- and Right-Handed Carbon Nanotubes by DNA. Journal of the American Chemical Society 2016, 138 (51), 16677–16685.

(31) Lyu, M.; Meany, B.; Yang, J.; Li, Y.; Zheng, M. Toward Complete Resolution of DNA/Carbon Nanotube Hybrids by Aqueous Two-Phase Systems. Journal of the American Chemical Society 2019, 141 (51), 20177–20186.

(32) Fagan, J. A. Aqueous two-polymer phase extraction of single-wall carbon nanotubes using surfactants. Nanoscale Advances 2019, 1 (9), 3307–3324.

(33) Yang, Y.; Zheng, M.; Jagota, A. Learning to predict single-wall carbon nanotube-recognition DNA sequences. npj Computational Materials 2019, 5 (1), 3.

(34) Li, H.; Gordeev, G.; Garrity, O.; Peyyety, N. A.; Selvasundaram, P. B.; Dehm, S.; Krupke, R.; Cambré, S.; Wenseleers, W.; Reich, S.; Zheng, M.; Fagan, J. A.; Flavel, B. S. Separation of Specific Single-Enantiomer Single-Wall Carbon Nanotubes in the Large-Diameter Regime. ACS Nano 2020, 14 (1), 948–963.

(35) Nißler, R.; Müller, A. T.; Dohrman, F.; Kurth, L.; Li, H.; Cosio, E. G.; Flavel, B. S.; Giraldo, J. P.; Mithöfer, A.; Kruss, S. Detection and Imaging of the Plant Pathogen Response by Near-Infrared Fluorescent Polyphenol Sensors. Angewandte Chemie International Edition 2022, 61 (2), e202108373.

(36) Nißler, R.; Bader, O.; Dohmen, M.; Walter, S. G.; Noll, C.; Selvaggio, G.; Groß, U.; Kruss, S. Remote near infrared identification of pathogens with multiplexed nanosensors. Nature Communications 2020, 11 (1), 5995.

(37) Spreinat, A.; Dohmen, M. M.; Lüttgens, J.; Herrmann, N.; Klepzig, L. F.; Nißler, R.; Weber, S.; Mann, F. A.; Lauth, J.; Kruss, S. Quantum Defects in Fluorescent Carbon Nanotubes for Sensing and Mechanistic Studies. The Journal of Physical Chemistry C 2021, 125 (33), 18341–18351.

(38) Harvey, J. D.; Williams, R. M.; Tully, K. M.; Baker, H. A.; Shamay, Y.; Heller, D. A. An in Vivo Nanosensor Measures Compartmental Doxorubicin Exposure. Nano Lett. 2019, 19 (7), 4343–4354.

(39) Galassi, T. V.; Jena, P. V.; Shah, J.; Ao, G.; Molitor, E.; Bram, Y.; Frankel, A.; Park, J.; Jessurun, J.; Ory, D. S.; Haimovitz-Friedman, A.; Roxbury, D.; Mittal, J.; Zheng, M.; Schwartz, R. E.; Heller, D. A. An optical nanoreporter of endolysosomal lipid accumulation reveals enduring effects of diet on hepatic macrophages in vivo. Science Translational Medicine 2018, 10 (461), eaar2680.

(40) Harvey, J. D.; Baker, H. A.; Mercer, E.; Budhathoki-Uprety, J.; Heller, D. A. Control of Carbon Nanotube Solvatochromic Response to Chemotherapeutic Agents. ACS Applied Materials & Interfaces 2017, 9 (43), 37947–37953.

(41) De los Santos, Z. A.; Lin, Z.; Zheng, M. Optical Detection of Stereoselective Interactions with DNA-Wrapped Single-Wall Carbon Nanotubes. Journal of the American Chemical Society 2021, 143 (49), 20628–20632.

(42) Turner, M. D.; Nedjai, B.; Hurst, T.; Pennington, D. J. Cytokines and chemokines: At the crossroads of cell signalling and inflammatory disease. Biochimica et Biophysica Acta (BBA) - Molecular Cell Research 2014, 1843 (11), 2563–2582.

(43) Kany, S.; Vollrath, J. T.; Relja, B. Cytokines in Inflammatory Disease. International Journal of Molecular Sciences 2019, 20 (23), 6008.

(44) Tertiş, M.; Ciui, B.; Suciu, M.; Săndulescu, R.; Cristea, C. Label-free electrochemical aptasensor based on gold and polypyrrole nanoparticles for interleukin 6 detection. Electrochimica Acta 2017, 258, 1208–1218.

(45) Cohen, Z.; Parveen, S.; Williams, R. M. Optimization of ssDNA-SWCNT Ultracentrifugation via Efficacy Measurements. ECS J. Solid State Sci. Technol. 2022, 11 (10), 101009.

(46) Gaelle, A.; Nathalie De, G.; Rino, M. Incorporation of Primary Amines via Plasma Technology on Biomaterials. In Advances in Bioengineering, Pier Andrea, S. Ed.; IntechOpen, 2015; p Ch. 2.

(47) Sims, C. M.; Zheng, M.; Fagan, J. A. Single-wall carbon nanotube separations via aqueous two-phase extraction: new prospects enabled by high-throughput methods. Chemical Communications 2025.

(48) Said, E. A.; Al-Reesi, I.; Al-Shizawi, N.; Jaju, S.; Al-Balushi, M. S.; Koh, C. Y.; Al-Jabri, A. A.; Jeyaseelan, L. Defining IL-6 levels in healthy individuals: A meta-analysis. J Med Virol 2021, 93 (6), 3915–3924.

(49) Góra-Gębka, M.; Liberek, A.; Szydłowskaysiak-Łysiak, W.; Bako, W.; Korzon, M. Serum interleukin 6 and interleukin 12 levels in children with chronic hepatitis HBV treated with interferon alpha. Annals of Hepatology 2003, 2 (2), 92–97.

(50) McCrae, L. E.; Ting, W.-T.; Howlader, M. M. R. Advancing electrochemical biosensors for interleukin-6 detection. Biosensors and Bioelectronics: X 2023, 13, 100288.

(51) Strand, V.; Boklage, S. H.; Kimura, T.; Joly, F.; Boyapati, A.; Msihid, J. High levels of interleukin-6 in patients with rheumatoid arthritis are associated with greater improvements in health-related quality of life for sarilumab compared with adalimumab. Arthritis Research & Therapy 2020, 22 (1), 250.

(52) Nicoletti, F.; Patti, F.; Cocuzza, C.; Zaccone, P.; Nicoletti, A.; Di Marco, R.; Reggio, A. Elevated serum levels of interleukin-12 in chronic progressive multiple sclerosis. Journal of Neuroimmunology 1996, 70 (1), 87–90.

(53) Chaiyaroj, S. C.; Rutta, A. S. M.; Muenthaisong, K.; Watkins, P.; Na Ubol, M.; Looareesuwan, S. Reduced levels of transforming growth factor-β1, interleukin-12 and increased migration inhibitory factor are associated with severe malaria. Acta Tropica 2004, 89 (3), 319–327.

(54) Frimpong, A.; Owusu, E. D. A.; Amponsah, J. A.; Obeng-Aboagye, E.; van der Puije, W.; Frempong, A. F.; Kusi, K. A.; Ofori, M. F. Cytokines as Potential Biomarkers for Differential Diagnosis of Sepsis and Other Non-Septic Disease Conditions. Front Cell Infect Microbiol 2022, 12, 901433.

(55) Nahar, Z.; Sal-Sabil, N.; Sohan, M.; Qusar, M. S.; Islam, M. R. Higher serum interleukin-12 levels are associated with the pathophysiology of major depressive disorder: A case-control study results. Health Sci Rep 2023, 6 (1), e1005.

(56) Stefan-van Staden, R.-I.; Ilie-Mihai, R. M.; Gugoasa, L. A.; Bilasco, A.; Visan, C. A.; Streinu-Cercel, A. Molecular recognition of IL-8, IL-10, IL-12, and IL-15 in biological fluids using phthalocyanine-based stochastic sensors. Analytical and Bioanalytical Chemistry 2018, 410 (29), 7723–7737.

(57) Bhavsar, K.; Fairchild, A.; Alonas, E.; Bishop, D. K.; La Belle, J. T.; Sweeney, J.; Alford, T. L.; Joshi, L. A cytokine immunosensor for Multiple Sclerosis detection based upon label-free electrochemical impedance spectroscopy using electroplated printed circuit board electrodes. Biosensors and Bioelectronics 2009, 25 (2), 506–509.

(58) Yang, T.; Wang, S.; Jin, H.; Bao, W.; Huang, S.; Wang, J. An electrochemical impedance sensor for the label-free ultrasensitive detection of interleukin-6 antigen. Sensors and Actuators B: Chemical 2013, 178, 310–315.

(59) Majdinasab, M.; Lamy de la Chapelle, M.; Marty, J. L. Recent Progresses in Optical Biosensors for Interleukin 6 Detection. Biosensors (Basel) 2023, 13 (9).

(60) Szymanska, B.; Lukaszewski, Z.; Oldak, L.; Zelazowska-Rutkowska, B.; Hermanowicz-Szamatowicz, K.; Gorodkiewicz, E. Two Biosensors for the Determination of Interleukin-6 in Blood Plasma by Array SPRi. Biosensors 2022, 12 (6), 412.

(61) Gordón, J.; Arruza, L.; Ibáñez, M. D.; Moreno-Guzmán, M.; López, M. Á.; Escarpa, A. On the Move-Sensitive Fluorescent Aptassay on Board Catalytic Micromotors for the Determination of Interleukin-6 in Ultra-Low Serum Volumes for Neonatal Sepsis Diagnostics. ACS Sensors 2022, 7 (10), 3144–3152.

(62) Muhammad, M.; Shao, C.-s.; Huang, Q. Aptamer-functionalized Au nanoparticles array as the effective SERS biosensor for label-free detection of interleukin-6 in serum. Sensors and Actuators B: Chemical 2021, 334, 129607.

(63) Jin, X.; Lee, M. A.; Gong, X.; Koman, V. B.; Lundberg, D. J.; Wang, S.; Bakh, N. A.; Park, M.; Dong, J. I.; Kozawa, D.; Cho, S.-Y.; Strano, M. S. Corona Phase Molecular Recognition of the Interleukin-6 (IL-6) Family of Cytokines Using nIR Fluorescent Single-Walled Carbon Nanotubes. Acs Appl Nano Mater 2023, 6 (11), 9791–9804.

