## Supplementary Figures File for "Multiplexed Molecularly-Specific SWCNT Cytokine Sensing is Enabled and Enhanced by Amine-Functionalized DNA Aqueous Two-Phase Extraction"

**Supplementary Figures S1-S7**

**Supplementary Tables S1-S2**

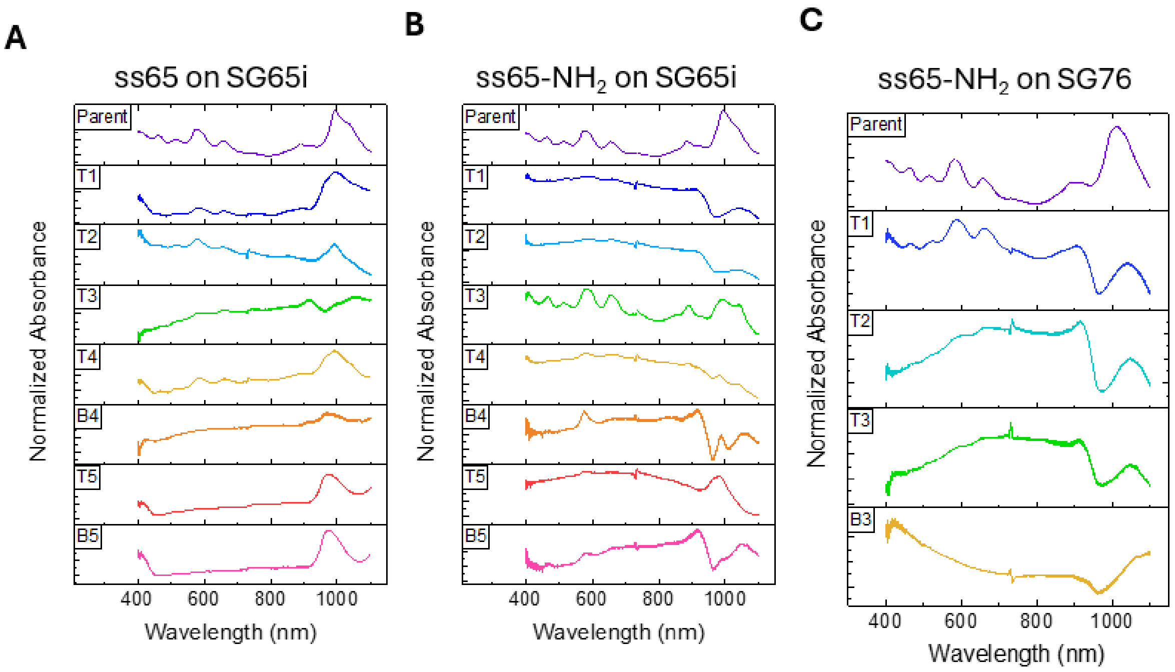


**Figure S1:** Absorbance spectra from each phase of ATPE sorting. **A)** SG65i SWCNT ss65. **B)** SG65i SWCNT using ss65-$NH_2$. **C)** SG76 SWCNT using ss65-$NH_2$.

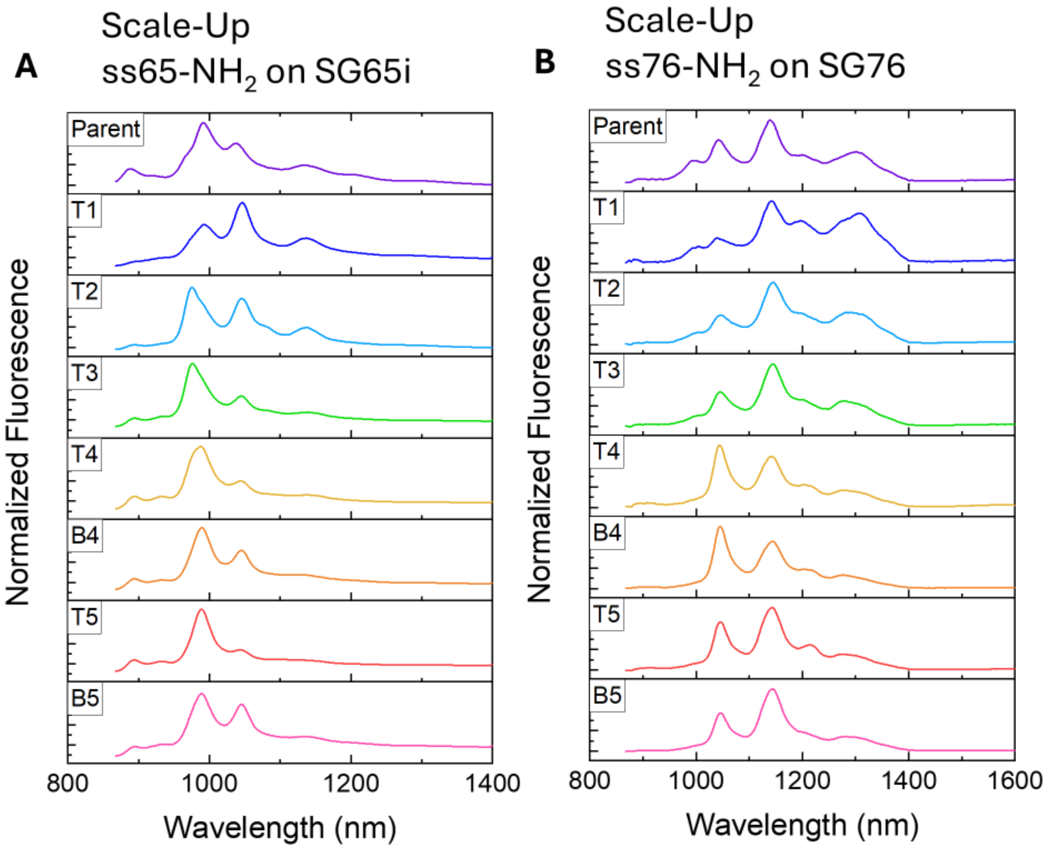


**Figure S2: Scale-up of ATPE sorting of SWCNT-DNA-$NH_2$. A)** Fluorescence spectra from each phase of ATPE sorting of SG65i using ss65-$NH_2$. **B)** Fluorescence spectra from each phase of ATPE sorting of SG76 using ss76-$NH_2$ (excitation wavelength = 638 nm).

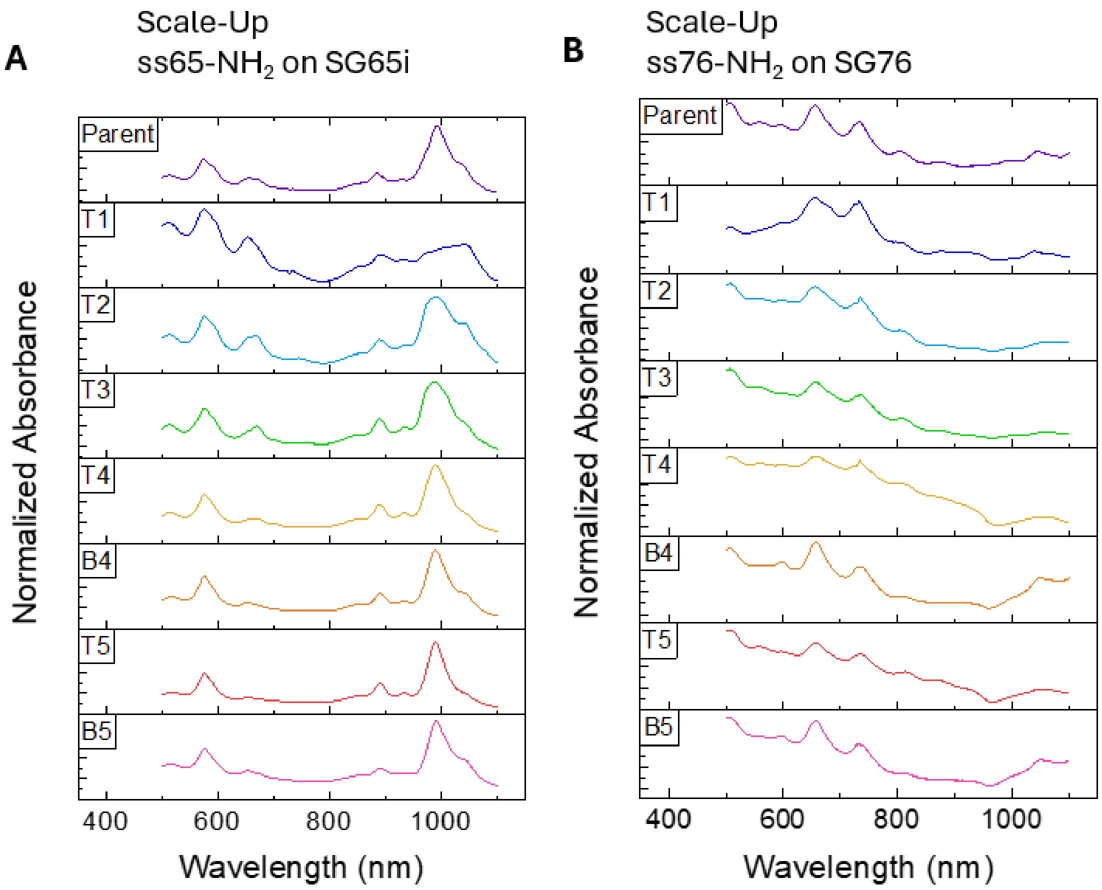


**Figure S3: Scale-up of ATPE sorting of SWCNT-DNA-$NH_2$. A)** Absorbance spectra from each phase of ATPE sorting of SG65i using ss65-$NH_2$. **B)** Absorbance spectra from each phase of ATPE sorting of SG76 using ss76-$NH_2$.

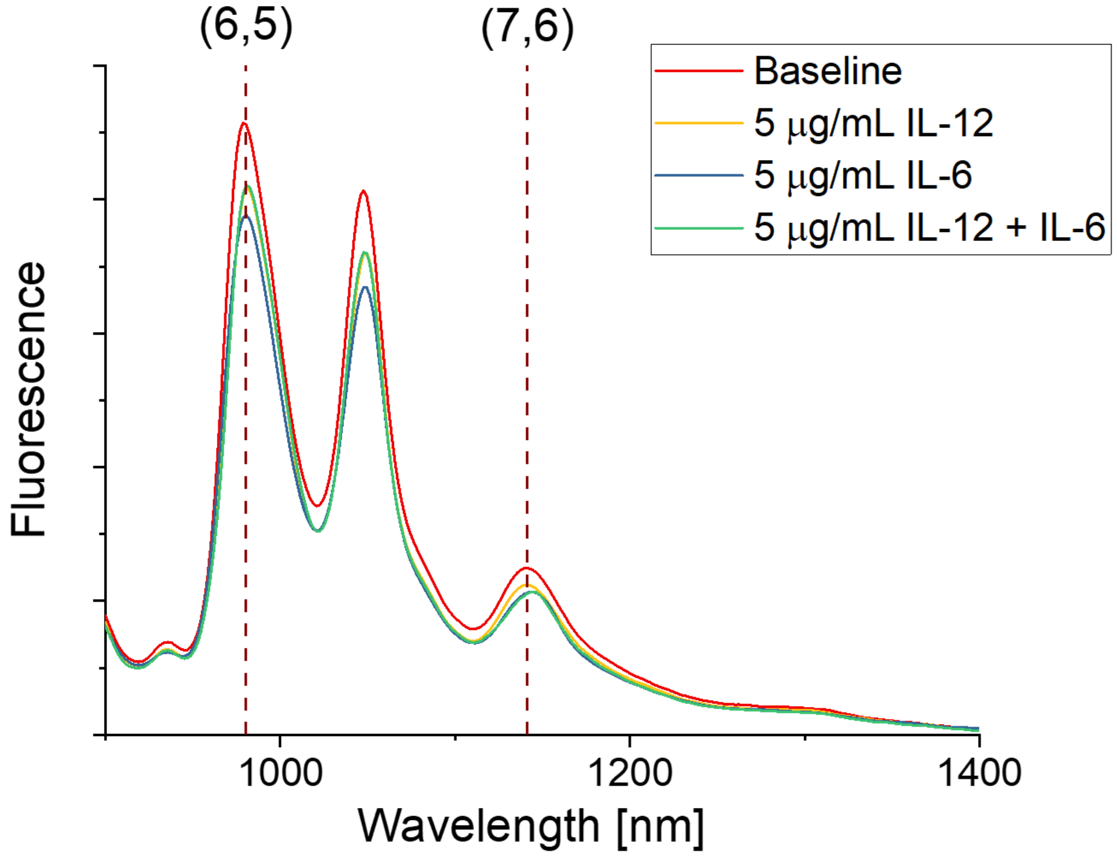


**Figure S4: NIR fluorescence spectra of individual and multiplexed cytokine detection (acquired with 655 nm laser)**

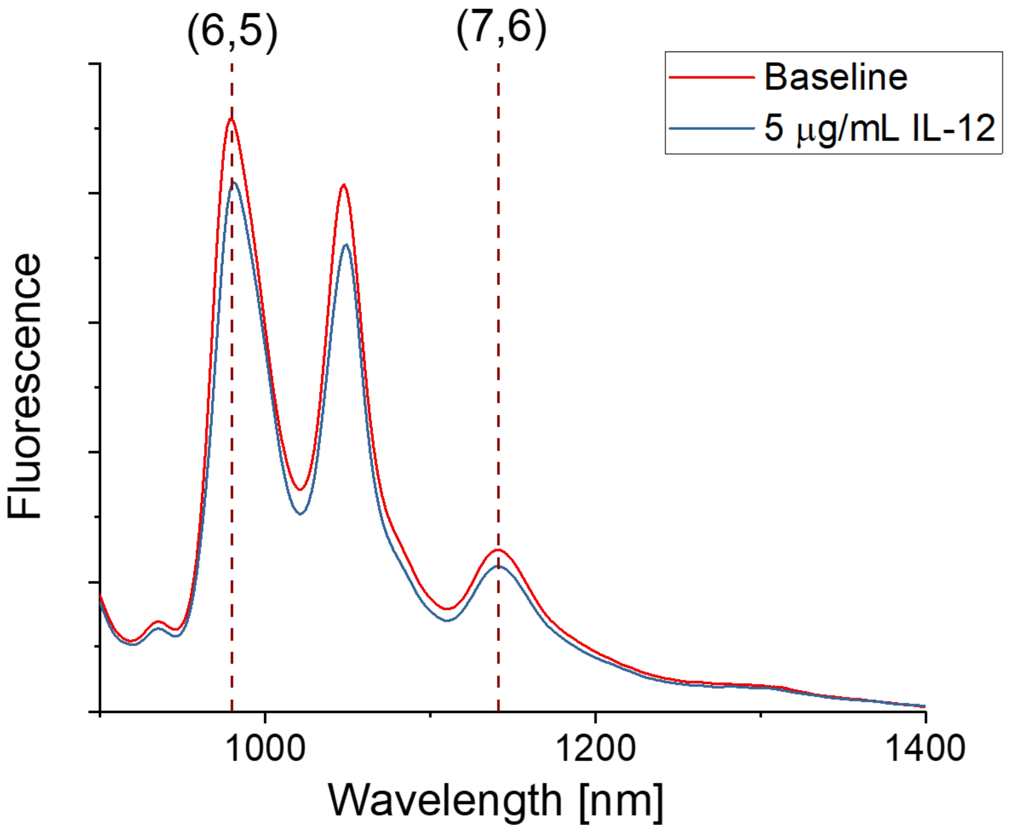


**Figure S5: NIR fluorescence spectra of IL-12 detection (acquired with 655 nm laser)**

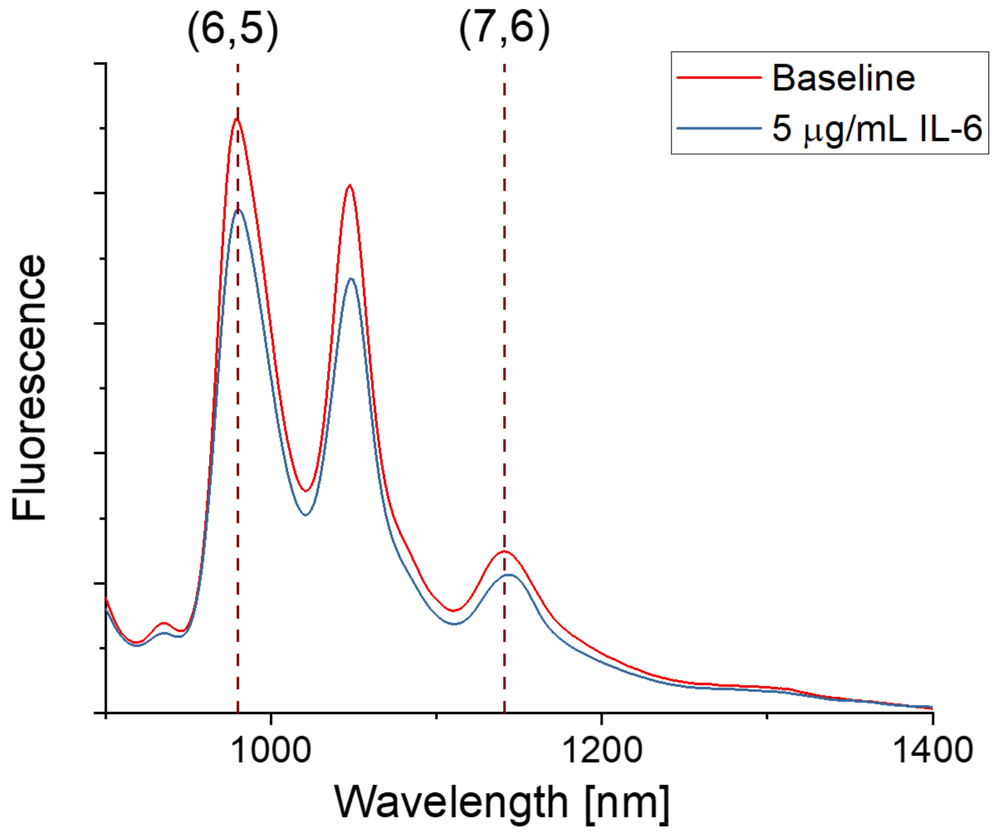


**Figure S6: NIR fluorescence spectra of IL-6 detection (acquired with 655 nm laser)**

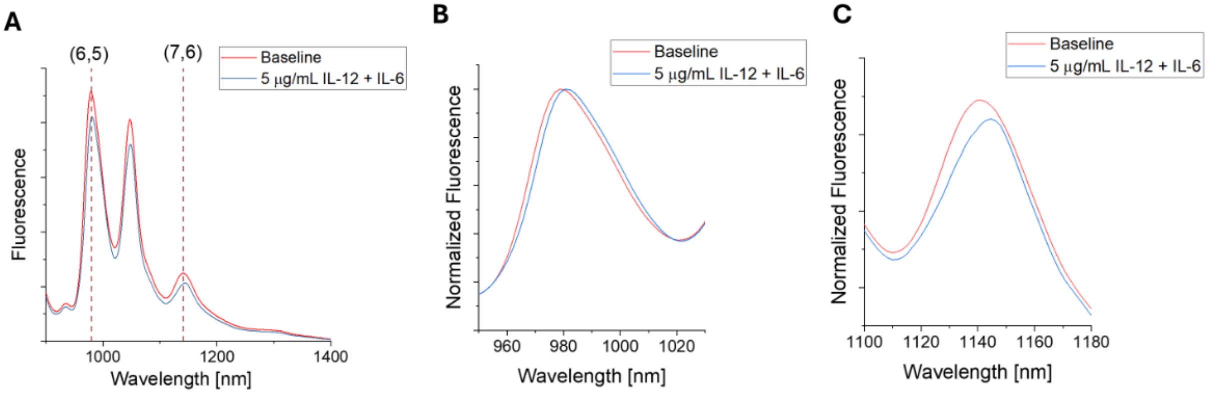


**Figure S7: NIR fluorescence spectra of simultaneous IL-12 and IL-6 detection. A)** Full NIR spectrum (acquired with 655 nm laser). B) (6,5) peak spectral shift. C) (7,6) peak spectral shift.

**Supplementary Table 1: (6,5) concentration-response curve p values**

| Cytokine Concentration (pg/mL) | IL-6 (denoted by *) | IL-12 (denoted by †) | IL6 + IL-12 (denoted by ‡) |
|---|---|---|---|
| 1.05E7 | 0.4749 | 1.16249E-4 | 2.82828E-5 |
| 5000000 | 0.0155 | 2.857E-4 | 0.00274 |
| 1500000 | 0.5484 | 5.674E-5 | 8.933E-5 |
| 500000 | 0.5861 | 9.838E-4 | 7.901E-5 |
| 50000 | 0.5887 | 2.057E-4 | 1.916E-4 |
| 15000 | 0.2107 | 8.402E-4 | 4.592E-4 |
| 5000 | 0.0949 | 2.965E-4 | 9.284E-4 |
| 1500 | 0.6815 | 7.2778E-4 | 5.319E-4 |
| 150 | 0.8251 | 7.308E-4 | 0.00176 |
| 15 | 0.693 | 0.00744 | 0.00358 |

**Supplementary Table 2: (7,6) concentration-response curve p values**

| **Cytokine Concentration (pg/mL)** | **IL-6 (denoted by *)** | **IL-12 (denoted by †)** | **IL6 + IL-12 (denoted by ‡)** |
|---|---|---|---|
| 1.05E7 | 1.75642E-6 | 0.00872 | 4.44266E-5 |
| 5000000 | 3.607E-5 | 0.02102 | 9.665E-6 |
| 1500000 | 2.0618E-5 | 0.63889 | 3.967E-5 |
| 500000 | 0.00313 | 0.00331 | 1.945E-4 |
| 50000 | 1.609E-5 | 0.02241 | 1.692E-5 |
| 15000 | 3.375E-5 | 0.80408 | 4.220E-5 |
| 5000 | 2.468E-4 | 0.35082 | 6.509E-6 |
| 1500 | 2.792E-4 | 0.77577 | 2.671E-4 |
| 150 | 5.298E-4 | 0.14737 | 9.998E-4 |
| 15 | 9.858E-4 | 0.23596 | 7.024E-4 |
